# A Combinatorial mRNA Therapy for Treating Rheumatoid Arthritis and Osteoarthritis by Inhibiting Inflammation and Promoting Cartilage Regeneration

**DOI:** 10.64898/2025.12.13.694161

**Authors:** Guan Wang, Yixuan Guo, Kun Guo, Cheng Zhang, Ying Zhou, Jiayuan Xia, Jiaqing Liu, Jinyi Ren, Ahmed Osman Mohamed, Ke Tang, Xianmei Chen, Shiyu Sun, Yong Yang, Mingyun Shen, Yifan Huang, Xia Li, Ang Lin

## Abstract

Rheumatoid arthritis (RA) and osteoarthritis (OA) are debilitating joint disorders with distinct etiologies but share common pathological features of chronic inflammation and progressive cartilage damage. Current therapeutic options are largely palliative and yet fail to achieve desired effect. In this study, we developed a combinatorial mRNA therapy using a rationally engineered lipid nanoparticle (LNP) containing a novel ionizable lipid for enhanced cartilage penetration upon intra-articular administration. This LNP co-delivers mRNAs encoding two complementary therapeutic proteins: interleukin-1 receptor antagonist (IL-1Ra) for inflammation attenuation and a C-terminal truncated derivative of angiopoietin-like 3 (ANL3) for cartilage regeneration. In pre-clinical models including collagen-induced arthritis mice, TNF-transgenic mice with spontaneous RA and destabilization of medial meniscus (DMM) surgical OA model, this “one-shot” combinatorial therapy significantly ameliorated disease severity, reduced synovitis and bone erosion, and potently promoted the regeneration of hyaline-like cartilage. Mechanistically, transcriptomic profiling of joint specimens revealed that the combinatorial therapy concurrently suppressed inflammatory pathways and extracellular matrix degradation pathways, meanwhile upregulating anabolic genes facilitating chondrogenesis. Collectively, our study establishes a versatile and synergistic mRNA therapy that offers a potent disease-modifying immunomodulatory strategy capable of addressing the long-standing unmet needs in arthritis treatment.

## Introduction

Arthritis, which is marked by severe joint dysfunction, impacts approximately 7–8% of the global population, with rheumatoid arthritis (RA) and osteoarthritis (OA) being the most prevalent forms(*1, 2*). RA is classified as an autoimmune disease, while OA is typically recognized as a degenerative joint disease. Despite their distinct pathogenesis, these two pathological conditions share significant similarities in multiple aspects, particularly regarding the cartilage damage and degree of inflammation(*3, 4*).

Therapeutic interventions for RA and OA differ significantly due to their distinct etiologies. Current approaches for RA treatment include disease-modifying antirheumatic drugs (DMARDs) and non-steroidal anti-inflammatory drugs (NSAIDs). These medications can to some degree alleviate disease activity by suppressing immune system and minimizing joint inflammation, but a subset of patients fail to respond adequately, and adverse effects remain a major concern(*5, 6*). While for OA treatment, it remains a significant challenge due to the lack of effective strategies. NSAIDs and exercise therapy are commonly employed for early-stage OA patients to slow disease progression, while surgical interventions such as joint replacement are often recommended for patients at advanced stage(*7, 8*). These treatments impose a significant burden on patients since no cure is currently available to reverse the joint damage, which underscores the urgent need for novel or improved therapies for arthritis.

Effective management of arthritis relies on two primary therapeutic strategies: reducing inflammation and promoting cartilage repair. Chronic inflammation is a defining feature and the main driver of disease progression, causing cartilage matrix breakdown, alterations in bone morphology, and the onset of pain(*4, 9*). A central mediator during this process is Interleukin-1 (IL-1), which upon binding to its receptor, initiates catabolic signaling that results in cartilage loss, synovitis, and bone erosion(*10*). A natural counteractor to IL-1 is the endogenous IL-1 receptor antagonist (IL-1Ra), which competitively inhibits IL-1 signaling by binding to its receptor without initiating the downstream inflammatory events(*11*). Indeed, a recombinant form of IL-1Ra, Anakinra, has been approved for RA treatment. However, its therapeutic efficacy is short-term and limited(*12*), likely due to the poor penetration into the cartilage matrix and its short half-life upon intra-articular administration.

Articular cartilage damage is another common feature of RA and OA, with both diseases involving the breakdown of cartilage matrix leading to progressive joint damage(*13*). Strategies to facilitate cartilage repair through the promotion of chondrocyte proliferation, differentiation, and extracellular matrix (ECM) synthesis may therefore hold promise in the management of arthritis. To address this, Novartis group utilized a high-throughput proteomic approach to discover the factors that facilitate chondrogenic differentiation of human mesenchymal stem cells. Angiopoietin-like 3 (ANGPTL3) was identified as a potent chondrogenic inducer, leading to the creation of its C-terminal truncated derivative with a K423Q mutation, known as LNA043(*14*). Pre-clinical evidence indicated that LNA043 could induce chondrogenesis, increase the production of lubricin, and regenerate hyaline articular cartilage(*14*). A first-in-human clinical trial demonstrated that LNA043 was well tolerated and could reverse OA transcriptional signature in OA patients^14^. These data positioned ANGPTL3 as a promising and effective target for therapeutic development against both RA and OA.

Despite the potential of emerging protein-based biologics for arthritis, limitations inherent to this approach persist in terms of short half-life, limited cartilage-penetrating capacity, and complex manufacturing procedure(*15*). Moreover, externally manufactured therapeutic proteins lack the post-translational modification profiles as naturally endogenous proteins, which may compromise their effectiveness. To tackle these challenges, leveraging mRNA platform for protein replacement therapy presents a more adaptable and revolutionary strategy that offers significant advantages over protein-based method, which has successfully been practiced in multiple disorders such as propionic acidaemia, fabry disease, psoriasis, etc(*16–18*). Nonetheless, the use of mRNA for protein replacement encounters a major hurdle, that is, precise delivery of mRNA cargo to affected tissues. The recently developed lipid nanoparticle (LNP) has become a prominent tool in mRNA field, which allows for precise modifications to the lipid component, molar ratio or nanoparticle size, enabling tailored biodistribution and optimal immunogenicity. While for articular cartilage delivery, the dense extracellular matrix presents the primary obstacle. In this regard, small nanoparticles (<100 nm, preferably <60 nm) have the advantage of enhanced cartilage matrix penetration upon intra-articular injection, as the mesh size of the collagen II fibrillar network is approximately 60 nm(*19–21*). To design an optimal LNP adaptable for cartilage delivery, we have developed a novel ionizable lipid FS01 that is constructed with a squaramide headgroup and an ortho-butylphenyl-modified hydrophobic tail(*17*), which can facilitate lipid-mRNA π-π interaction in favour of formulating LNP with a small diameter. Utilizing this unique LNP system, we strategically designed a combinatorial mRNA therapy encoding IL-1Ra and a modified form of ANGPTL3 (ANL3) to treat RA and OA by simultaneously inhibiting inflammation and promoting cartilage regeneration. In multiple mouse models, including collagen-induced arthritis (CIA) mice, TNF transgenic mice (TNF-Tg) with spontaneous RA, and destabilization of medial meniscus (DMM) surgical OA model, this dual mRNA therapy showed potent efficacy in treating RA and OA, as evidenced by reduced arthritis index, decreased swelling and joint inflammation. Furthermore, the expression of collagen II, an essential component of articular cartilage supporting joint function, was restored, thus indicating cartilage regeneration following treatment. Furthermore, RNA sequencing (RNA-seq) on joint samples collected from CIA mice demonstrated that this combinatorial therapy effectively attenuated inflammatory pathways and extracellular matrix degradation pathways, which elucidated the core mechanism of action.

## Materials and Methods

A detailed description and additional methods are available in supplemental materials.

### Ethics

All animal experiments were conducted according to the Guidelines for the Care and Use of Laboratory Animals and the Ethical Committee of China Pharmaceutical University and Dalian Medical University. Protocols were approved by the Institutional Animal Care and Use Committee of China Pharmaceutical University (Approval number: CPUTML-LA-20220109) and Dalian Medical University (Approval number: AEE24013). All animals were housed in a specific pathogen-free (SPF) facility where the temperature was maintained at 22 ± 1°C and a 12-hour light/dark cycle was used.

### RA and OA mouse models

For establishment of collagen-induced arthritis (CIA) mice, male DBA/1J mice, aged 6–8 weeks, were procured from Slack Laboratory Animal Co., Ltd (Shanghai, China) and acclimated for 1 week. To induce arthritis, mice received a subcutaneous (s.c) injection of a 100-μl emulsion containing bovine type II collagen (Chondrex, USA) mixed with complete Freund’s adjuvant (CFA, Chondrex, USA) at a 1: 1 volume ratio on day 0, followed by another s.c administration of a 100-μl emulsion containing bovine type II collagen mixed with incomplete Freund’s adjuvant (IFA, Chondrex, USA) at a 1: 1 volume ratio on day 21. For establishment of spontaneous RA mouse model, TNF-Tg mice aged 12 weeks with a C57BL/6 background were used, which developed a progressive, erosive arthritis accompanied by significant bone loss. For establishment of OA mouse model, experimental OA was induced by surgical DMM on C57BL/6 mice (male, 6 weeks old) according to our well-established protocol reported previously(*22*). Briefly, mice were anesthetized with isoflurane, and the right knee joint capsule was opened. The medial meniscotibial ligament, which secures the medial meniscus to the tibial plateau, was transected to destabilize the joint, while cartilage injury beneath the medial meniscus was avoided. Regarding the animals in sham control group, the knee joint capsule was opened but not transected before being closed.

### mRNA design and in vitro transcription (IVT)

IL-1Ra-mRNA and ANL3-mRNA were designed and codon-optimized using our proprietary algorithm, primarily based on the codon-adaptation index (CAI) and minimum free energy (MFE). The mRNA sequences incorporated the 5ʹ-UTR of human α-globin, AES-mtRNR1 3ʹ-UTR deployed in BNT162b2, and a 120 nt poly A tail, which were enlisted in Supplemental Table 1. mRNAs with purity exceeding 95% were synthesized using a well-established protocol as previously reported by our group(*17, 23–25*). In brief, mRNAs were synthesized via a T7 polymerase-mediated IVT procedure using a linearized DNA template (pUC57-GW-Kan) containing 5’-UTRs, 3’-UTRs and a 120-nt poly (A) tail. During the IVT step, mRNAs were chemically modified with N1-Methyl pseudo-uridine (NEB) and capped with cap analogues (LZCap AG, C_35_H_48_FN_15_O_24_P_4_ or TriLink CleanCap AG, C_32_H_43_N_15_O_24_P_4_). The mRNA products were then purified using Monarch RNA purification columns (NEB) and dissolved in TE buffer at a desired concentration for further applications.

### LNP formulation and characterization

LNPs were formulated using a well-established protocol as described previously(*17, 25*), by mixing an organic phase containing lipids with an aqueous phase containing mRNAs at a volume ratio of 3: 1 using a microfluidic-based equipment (INanoL from Micro&Nano Biologics) at a total flow rate of 12 mL/min. The organic phase was prepared by dissolving ionizable lipid, 1,2-distearoyl-sn-glycero-3-phosphocholine (DSPC), cholesterol, and DMG-PEG2000 (1,2-dimyristoyl-rac-glycero-3-methoxypolyethylene glycol-2000) in ethanol at molar ratios of 50: 10: 38.5: 1.5. Four different ionizable lipids were used in this study including SM-102, Lipid 5, LP01 and FS01. The ionizable lipid FS01 is a proprietary lipid that was synthesized in-house and has been well-characterized in our previous study(*17*). Other three ionizable lipids were purchased from XIAMEN SINPOEC BIOTECH. The aqueous phase was prepared in a 10 mM citrate buffer (pH 4.0) containing corresponding mRNA payloads (firefly luciferase-mRNA, IL-1Ra-mRNA, or ANL3-mRNA). Upon encapsulation, the mixture was diluted with PBS and ultrafiltrated using 50-kDa Amicon ultracentrifugal filters (Millipore). Encapsulation efficiency and mRNA concentration were determined using the Quant-iT™ RiboGreen RNA assay (Invitrogen, US). The hydrodynamic diameters, polydispersity index (PDI), and zeta potential of the LNPs were measured using BeNano 90 Zeta nanoparticle size and zeta potential analyzer (Dandong Bettersize, China).

### In-vivo bioluminescence study

To assess the intra-articular delivery efficiency of different LNPs, C57BL/6 mice (male, 6-week-old) were administered with Fluc-mRNA-LNPs (5 μg, n = 4/group) via joint cavity injection. 6, 12, 24, and 48 hours post injection, mice were intraperitoneally (i.p.) injected with 150 mg/kg D-luciferin (Beyotime, China), and bioluminescence images were captured using the In Vivo Imaging System (IVIS) Spectrum (Perkin Elmer, Santa Clara, CA) within 20 min. The exposure time was set to 10 seconds, and bioluminescence signals were quantified using the Living Image software (PerkinElmer).

### Treatment and evaluation

To evaluate the therapeutic effects in RA mouse model, mice received three or four doses of intra-articular injection with IL-Ra-mRNA-LNP (2 μg), ANL3-mRNA-LNP (2 μg), or IL-1Ra/ANL3-mRNA-LNP (4 μg) at an interval of 7 days. Mice administered with PBS were used as control. At the day of necropsy, hind paw diameters were measured by a digital caliper. The arthritis index was evaluated using the following criteria: 0, normal; 1, mild swelling and redness confined to the tarsals or ankle joint; 2, mild swelling and redness extending from the ankle to the tarsals; 3, moderate swelling and redness extending from the ankle to metatarsal joints; and 4, severe swelling and redness encompassing the ankle, foot and digits. The ankle joints specimens were harvested for micro-computed tomography (micro-CT) analysis. Images were collected and reconstructed using the 3D reconstruction software Recon. To evaluate the therapeutic effects in OA mouse model, mice were treated with two or three doses of IL-1Ra/ANL3-mRNA-LNP (4 μg) via intra-articular injection at an interval of 7 days. Body weight of animals was monitored weekly during the experiment. At the day of necropsy, joint specimens were collected and subjected to hematoxylin and eosin (H&E) staining and Safranin O-Fast Green (SO/FG) staining for assessment. Histopathological grading was evaluated by Mankin’s scoring (0 – 14 points) based on cartilage structure, cellularity, cartilage matrix and tidemark integrity(*26*). SO/FG staining was semi-quantitatively assessed by Osteoarthritis Research Society International (OARSI) scoring (0–6 points) based on the depth and extent of cartilage structure damage(*27*).

### Flow cytometric analysis

For surface staining, cells were stained with LIVE/DEAD™ Fixable Aqua Dead Cell Stain Kit (Thermo) for 5 minutes and then incubated with antibody cocktails and Fc receptor blocking reagent (Miltenyi) for 20 minutes at 4°C in dark. For intracellular nuclear staining, cells were first fixed and permeabilized using Fixation/Permeabilization Solution Kit (BD Biosciences), followed by staining with the nuclear antibody anti-mouse Foxp3-PE (eBioscience, USA) for 30 minutes. For intracellular cytokine staining, cells were pre-stimulated with phorbol myristate acetate (PMA) (50 ng/mL) and ionomycin (1 μg/mL) for 4 hours in the presence of brefeldin A (10 μg/mL) to block cytokine secretion. Following surface staining, cells were fixed, permeabilized, and then stained intracellularly with antibodies against cytokines including IFN-γ, IL-4, IL-17, or TNF. Data were acquired on a Flow Cytometer (Agilent, USA) and analyzed using NovoExpress software. A detailed list of antibodies used in flow cytometric analysis is available in Supplemental Table 2.

### Statistical analysis

Statistical analysis was performed with GraphPad Prism 9 software (version 9.4, San Diego, USA). Statistical differences were analyzed by unpaired t-test, Mann-Whitney test, or One-way ANOVA with post-hoc Tukey’s test for correction. Data are presented as mean ± Standard Error of the Mean (SEM). A *p* value less than 0.05 was considered statistically significant. Asterisks mark the significant differences between different groups. *\* p < 0.05; ** p < 0.01, *** p <0.001, **** p <0.0001*.

## Results

### Rational design of a dual mRNA therapy for RA and OA using a novel ionizable lipid-LNP favoring cartilage penetration

To develop a novel mRNA therapy for RA and OA, we for the first time, from the perspective of anti-inflammation and cartilage regeneration, selected IL-1Ra and a C-terminal truncated form of ANGPTL3 (abbreviated as ANL3) as the druggable targets. Utilizing a well-established mRNA-LNP platform(*23, 25, 28, 29*), we encapsulated IL-1Ra-mRNA and ANL3-mRNA mixed at an equivalent mass ratio into a unique LNP system, which was administered into the knee joint cavity for therapeutic purpose (Fig. 1A).

**Figure 1.**
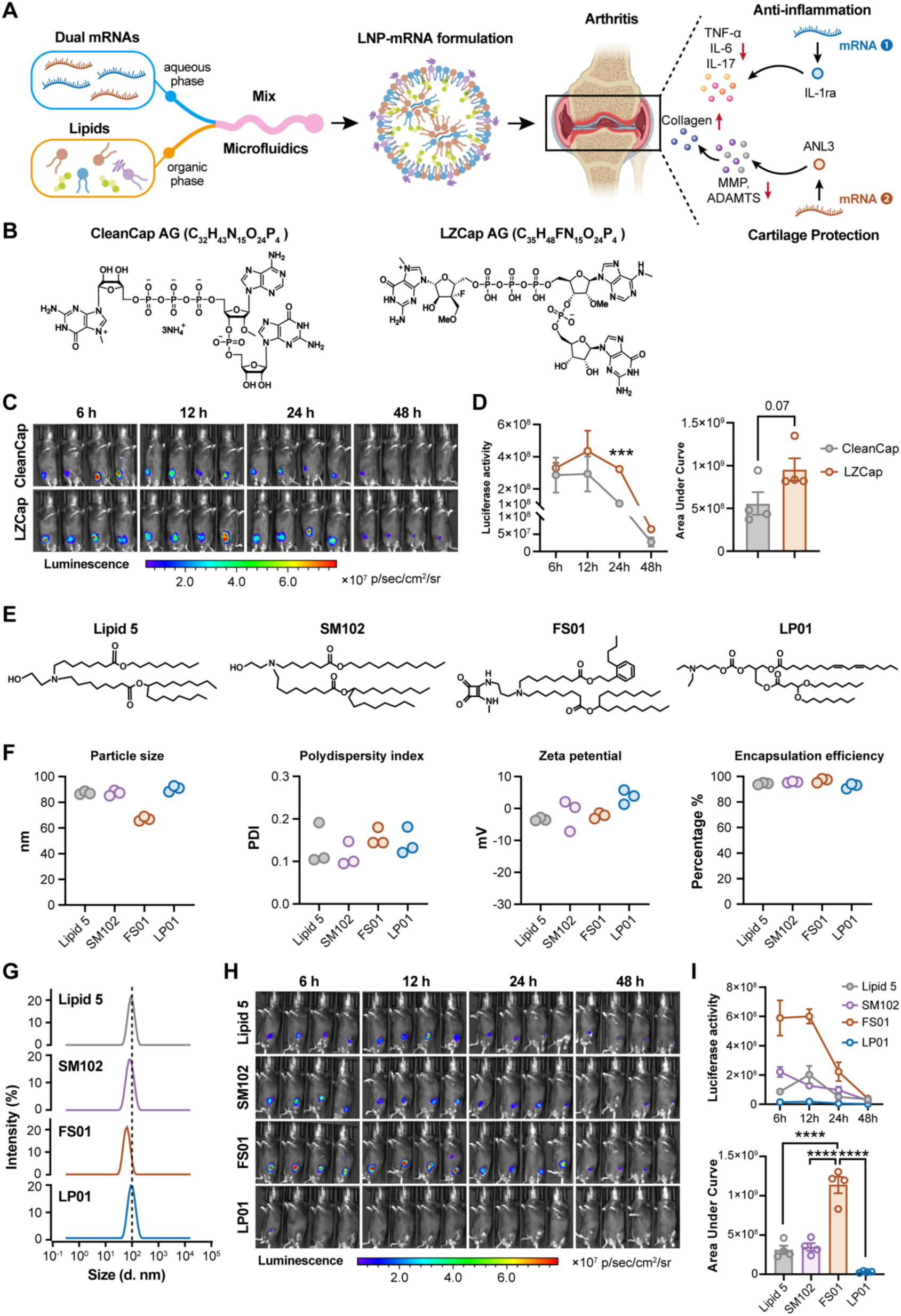
Rational design of LNP-mRNA for optimal delivery to the intra-articular cartilage compartment. **A,** Experimental design schematic. Two separate mRNAs encoding for target proteins (IL-1Ra and ANL3) are mixed at an equivalent mass ratio, followed by co-encapsulation into LNP. Upon intra-articular injection, IL-1Ra-mRNAs and ANL3-mRNAs are released and their correspondingly expressed proteins function to treat arthritis disease, including RA and OA. IL-1Ra reduces a cascade of inflammatory mediator and ANL3 promotes chondrogenesis and cartilage repair. **B,** Structure of two Cap analogs used for mRNA synthesis, CleanCap AG (C_32_H_43_N_15_O_24_P_4_) and LZCap AG (C_35_H_48_FN_15_O_24_P_4_). **C-D,** LNP-Fluc-mRNA constructed with different Cap analogs were tested for expression at local sites following intra-articular injection. **C,** Bioluminescent images (n=4/group) are shown. **D,** Quantification of luminescence at the indicated time point and total luminescence calculated as area under curve (AUC) are shown. **E,** Structures of ionizable cationic lipids, including Lipid 5, SM102, FS01 and LP01. **F,** Major parameters including particle size, PDI, zeta potential, and encapsulation efficiency of the four LNP formulations are indicated. **G,** Particle size of mRNA-LNP prepared with different ionizable lipids measured by dynamic light scattering. **H,** Delivery efficiency of Fluc-mRNA by different LNPs upon intra-articular injection was evaluated. Bioluminescent images (n=4/group) are shown. **I,** Quantification of luminescence at the indicated time point and total luminescence calculated as AUC are shown. Data are shown as mean ± SEM. One-way ANOVA test or unpaired t test was used for statistical analysis. *** *p* <0.001; **** *p* <0.0001.

During mRNA synthesis, N1-methyl-pseudouridine (m1Ψ) was incorporated to reduce mRNA’s intrinsic immunogenicity and improve stability. In addition, the Cap structure at the 5’ end of mRNA plays a critical role in regulating mRNA translation and stability(*30*). Among various Cap analogs used in mRNA industry, the Cap 1 analog, CleanCap AG (C_32_H_43_N_15_O_24_P_4_), is commonly employed during mRNA synthesis through a co-transcriptional capping approach(*31*). While in our study, to further enhance the translational efficiency of mRNA, we utilized a novel proprietary Cap 1 analog LZCap AG (C_35_H_48_FN_15_O_24_P_4_) to construct the mRNAs (Fig. 1B). Firefly luciferase (Fluc)-mRNAs constructed with the two different Cap 1 analogs were encapsulated into LNP and administered via intra-articular route into C57BL/6 mice for evaluation of their *in-vivo* translation. Bioluminescent signals were longitudinally detected at 6, 12, 24 and 48 hours post injection (Fig. 1C). Fluc-mRNA capped with LZCap AG demonstrated a more robust translation efficacy at the site of administration than the control mRNA (Fig. 1D).

Delivering therapeutic mRNA via LNP into the affected intra-articular cartilage compartment remains challenging due to that conventional LNPs with diameters between 100–200 nm have a very limited capacity to penetrate the dense extracellular matrix layer. On the other hand, the ionizable lipid component within LNP is inherently endowed with inflammatory property(*32*), which should be minimized for protein replacement therapy, otherwise may compromise the efficacy and safety. Previously, we reported on a low-inflammatory ionizable lipid FS01 that can form small-sized LNP (around 60 nm) owning to its unique structure featured by a squaramide headgroup and an ortho-butylphenyl modified hydrophobic tail(*17*). To design an optimal LNP that benefits intra-articular mRNA delivery, we evaluated and compared FS01 with three other ionizable lipids including Lipid 5, SM102, and LP01 (Fig. 1E) that have been used in approved mRNA vaccine(*33*) or other mRNA therapies at clinical stages (Clinical trial nr. NCT04601051)(*34, 35*). Fluc-mRNA-LNP was formulated using a classical formulation containing ionizable lipid, cholesterol, DSPC, and PEG-DMG at molar ratios of 50: 10: 38.5: 1.5%. Notably, FS01-LNP displayed a smaller particle size than other three LNPs, achieving an average diameter of 67 nm (Fig. 1, F and G). All the four LNPs showed a polydispersity index (PDI) between 0.1 and 0.2, a zeta-potential value between - 10 and + 10 mV, and a high encapsulation efficiency exceeding 90% (Fig. 1F and fig. S1). Further, upon injection into the knee joint cavity, kinetics of bioluminescent signals derived from mRNA translation and accumulative signals during the period of observation were calculated. FS01-LNP was significantly more efficient than other LNPs in delivering mRNA into the articular compartment, while interestingly, LP01-LNP showed no or very limited mRNA delivery capacity (Fig. 6, H and I). These data supported that FS01-LNP is a rationally constructed system adaptable for intra-articular delivery of mRNA cargo.

Using the design strategies above, IL-1Ra-mRNA and ANL3-mRNA were constructed and IVT synthesized. To facilitate secretion of translated proteins, a signal peptide of human immunoglobulin kappa chain (Ig κ) was inserted into the 5’-end of sequence. Upon transfection into HEK-293T cells for 24 hours, IL-1Ra-mRNA and ANL3-mRNA were efficiently translated, and the corresponding proteins were readily detectable in culture supernatants, as measured by ELISA and Western blot assay, respectively (Fig. 2, A and B). We then encapsulated IL-1Ra-mRNA and ANL3-mRNA either separately or in a mixed form into the FS01-LNP. All major physiochemical parameters of mRNA-LNP formulations including particle size, PDI, zeta potential, encapsulation efficiency met the standards, confirming the high quality of products (Fig. 2C). In addition, to prove that FS01-LNP is capable to deliver mRNA into the cartilage site, we administered 2 μg of IL-1Ra-mRNA-LNP intra-articularly into C57BL/6 mice and detected a robust expression of IL-1Ra localized at the cartilage site (fig. S2). Kinetics and expression of IL-1Ra in situ derived from IL-1Ra-mRNA (2 μg) was compared side-by-side with that from directly administered recombinant IL-1Ra protein (2 μg). A significantly higher magnitude of protein expression was found in the IL-1Ra-mRNA group, which suggested the advantage of using mRNA for a more sustained and pronounced local protein production in this specific context (fig. S2).

**Figure 2.**
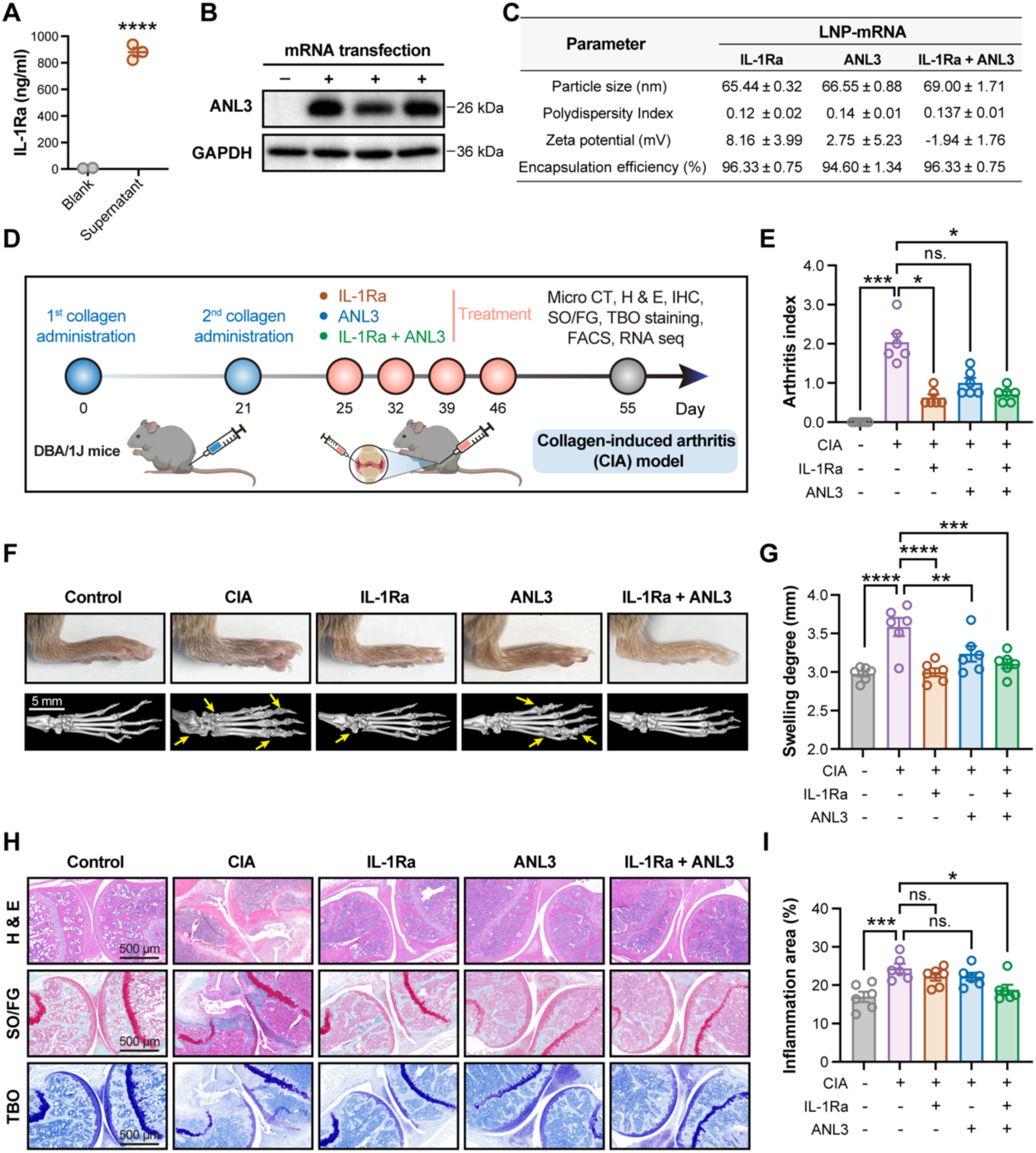
Therapeutic effect of mRNA therapies in CIA mouse model. **A-B,** Translation of IL-1Ra-mRNA and ANL3-mRNA upon transfection into HEK-293T cells for 24 hours. Protein levels of IL-1Ra (**A**) and ANL3 (**B**) in cell culture supernatants were quantified by ELISA and Western Blot, respectively. Data is from three independent experiments. **C,** A summary of major physiochemical parameters of mRNA-LNP formulations. **D,** Experimental design schematic. DBA/1J mice were immunized with collagen type II on days 0 and 21 for CIA induction. Mice were treated with 2 μg of IL-1Ra-mRNA-LNP, 2 μg of ANL3-mRNA-LNP, or 4 μg of IL-1Ra/ANL3-mRNA-LNP on day 25, 32, 39 and 46. At day 55, mice were necropsied for analyses. Healthy DBA/1J represented the healthy control (n=6 mice/group). **E,** Arthritis index is shown. **F,** Representative images from morphological appearance (upper panel) and micro-CT scanning (lower panel) of ankle joints and paws of mice. Arrows indicate sites of bone erosion. **G,** Paw thickness indicative of swelling degree (n=6 mice/group). **H,** Representative images of H&E, safranin O/fast green (SO/FG) and toluidine blue (TBO) staining of the tissue sections from knee joints. **I,** Inflammatory cell infiltration was quantified by measuring the percentage of inflammation-positive area using Image J. Data are displayed as the mean ± SEM. Statistical significance was determined using One-way ANOVA with Tukey’s post hoc test. * *p* < 0.05; ** *p* < 0.01; *** *p* < 0.001; **** *p* < 0.0001.

### IL-1Ra/ANL3-mRNA-LNP ameliorates RA progression in collagen-induced arthritis (CIA) mice

Next, we comprehensively evaluated the therapeutic potential of mRNA therapy in CIA mouse model. DBA/1J mice were primarily immunized with two doses of bovine type II collagen on day 0 and 21 for arthritis induction, and subsequently received four doses of monotherapy with IL-1Ra-mRNA-LNP (2 μg) or ANL3-mRNA-LNP (2 μg), or dual therapy with IL-1Ra/ANL3-mRNA-LNP (4 μg) at an interval of seven days (Fig. 2D). By day 55, mice treated with IL-1Ra-mRNA-LNP alone or with IL-1Ra/ANL3-mRNA-LNP dual therapy demonstrated a significantly slowed disease progression. This was directly evidenced by the reduced arthritis index (Fig. 2E), which represents the disease severity by scoring clinical signs like redness and swelling. ANL3-mRNA-LNP also showed a moderate therapeutic potential in terms of arthritis index reduction, although not significant (Fig. 2E). In addition, morphological observation revealed that all three groups of treated mice showed a dramatically lower degree of paw swelling compared to the untreated mice (Fig. 2, F and G). We further deployed a micro-CT imaging approach to assess articular bone erosion in the ankle joints and paws of mice. The untreated CIA mice showed irregular bone surfaces and severe bone erosion in the ankle and metatarsophalangeal joints. While in contrast, monotherapy with IL-1Ra-mRNA-LNP and dual therapy with IL-1Ra/ANL3-mRNA-LNP significantly reduced bone erosion degree, which was more prominently observed in the dual therapy group. ANL3-mRNA-LNP also ameliorated bone damage to some degree but was less efficient than other two treatments (Fig. 2F).

Histological examinations by H&E staining were also performed on the knee joint tissues. Compared to the untreated CIA mice, all three groups of treated mice exhibited significantly improved cartilage integrity and reduced synovitis accompanied by a lower degree of inflammatory cell infiltration into the synovium. Importantly, the IL-1Ra/ANL3-mRNA-LNP treatment induced a more pronounced effect than the two monotherapy groups (Fig. 2H, upper panel). While of note, treatment with ANL3-mRNA-LNP alone or the dual therapy led to a noticeable increase in cartilage thickness, as revealed by safranin O/fast green (SO/FG) and toluidine blue (TBO) staining that delineated the degree of articular cartilage degradation (Fig. 2H), which was not observed in the IL-1Ra-mRNA-LNP treatment group. These data suggested a critical role of ANL3 and IL-1Ra in promoting cartilage regeneration and inhibiting inflammation, respectively, which supported the rationale behind drug development by deploying two distinct mechanisms for arthritis treatment synergistically.

### Transcriptomic analysis reveals modulation of inflammatory and extracellular matrix degradation pathways in CIA mice upon mRNA therapy

To further clarify therapeutic efficacy and gain more insights into the mechanism of action, we conducted RNA sequencing (RNA-seq) and transcriptomic analysis on the knee joint specimens. Principal component analysis (PCA) revealed distinct gene expression profiles of the control naïve mice, untreated and mRNA-treated CIA mice (Fig. 3A). As expected, a substantial number of differentially expressed genes (DEGs) were identified when comparing the untreated CIA mice to the healthy control. While pairwise comparisons between each treatment group and the untreated group yielded a substantially smaller number of DEGs (Fig. 3B). To identify the core pathways altered by mRNA treatments, we pooled the DEGs obtained from all “treatments *vs.* CIA” comparisons and selected 185 genes for Gene Ontology (GO) and Kyoto Encyclopedia of Genes and Genomes (KEGG) pathway enrichment analyses (Fig. 3C). The results indicated that the DEGs were significantly enriched in pathways related to inflammation and extracellular matrix (ECM) degradation, aligning closely with the mechanisms of IL-1Ra and ANL3 (Fig. 3C).

**Figure 3.**
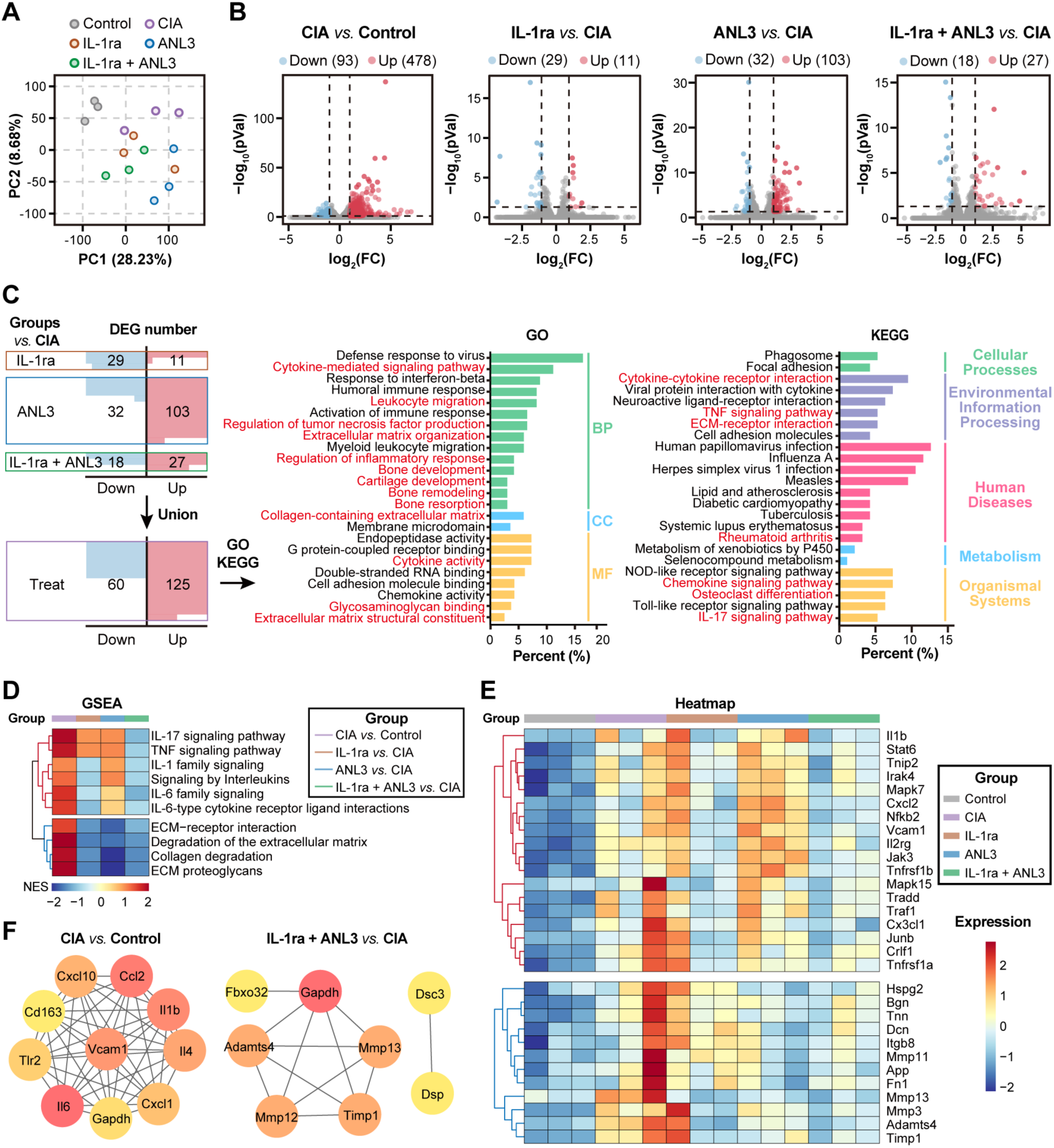
IL-1Ra/ANL3-mRNA-LNP modulated inflammatory and extracellular matrix degradation pathways in CIA mice. CIA mice were treated following the experimental schedule shown in Figure 2b. RNA samples of knee joint specimens from two mice per group were pooled for RNA sequencing (n=3 biological replicates in each group). **A,** Principal component analysis (PCA) of genes. **B,** Volcano plots illustrating DEGs for each treatment group compared to the untreated CIA group. Criteria used are *p* value < 0.05 calculated using a paired two-tailed Student’s *t* test and at least 2-fold change following treatment. **C,** GO and KEGG enrichment analyses on the total DEGs across all treatment groups. **D,** Heatmap of Gene Set Enrichment Analysis (GSEA) results for modules related to inflammation pathways and bone destruction and formation. **E,** Heatmap indicates the expression levels of representative genes selected from the modules in panel d. **F,** Protein-protein interaction (PPI) network of DEGs, constructed using data from the STRING database.

We next performed Gene Set Enrichment Analysis (GSEA) to dissect the detailed molecular mechanisms underlying the therapeutic effects of IL-1Ra and ANL3 mRNA-LNPs. Compared to healthy mice in control group, CIA mice exhibited a marked upregulation of genes associated with inflammation and ECM degradation. IL-1Ra-mRNA-LNP monotherapy primarily downregulated genes associated with IL-1 and IL-6 inflammatory signaling and concurrently reduced the ECM pathways (Fig. 3D). Conversely, ANL3-mRNA-LNP monotherapy had minimal impact on inflammatory genes but significantly suppressed the ECM degradation pathways. Notably, the dual mRNA therapy effectively attenuated both inflammatory pathways and ECM pathways (Fig. 3D), which indicated a synergistic effect between IL-1R-mRNA and ANL3-mRNA therapy. Representative genes associated with inflammation such as *il1b, cxcl2, nfkb2* and genes related to ECM degradation such as *mmp11, mmp 13, adamts4* were differentially altered by different mRNA treatments (Fig. 3E), which was in line with the GSEA analysis. Finally, we performed a Protein-Protein Interaction (PPI) network analysis on the DEGs. In CIA mice, PPI network was dominated by hubs associated with inflammation and chemotaxis, underscoring the pathological drivers of the arthritis diseases. In contrast, upon mRNA therapy, the network architecture shifted, and key modules became enriched with proteins involved in matrix degradation (Fig. 3F). Overall, the transcriptomic analysis revealed a core mechanism of action for the IL-1Ra/ANL3-mRNA therapy through concurrent suppression of inflammatory cascades and ECM degradation pathways.

### IL-1Ra/ANL3-mRNA-LNP therapy inhibits inflammation and promotes cartilage regeneration in CIA mice

To validate the findings from RNA-seq analysis, we next performed a comprehensive immunological assessment. Th1 and Th17 cells are two major pro-inflammatory cell subsets that have been extensively reported to contribute to the pathogenesis of RA(*36*). One typical Th1-type cytokine, TNF-α, has been shown to induce a pro-inflammatory microenvironment in the synovium that leads to cartilage destruction and bone erosion(*37*). IL-17A released by Th17 cells is also highly involved in the immune regulation of RA by not only over-activating inflammatory cascades but also promoting osteoclast differentiation leading to cartilage damage(*38, 39*). Moreover, TNF-α and IL-17A tightly crosstalk with each other in a synergistic way to amplify the inflammatory signals favoring RA progression(*40*). Flow cytometry analysis of splenic CD4^+^ T cells revealed that all mRNA-LNP treatments significantly reduced the production of TNF-α and IL-17A by T cells (Fig. 4A). Consistent with this, levels of serum TNF-α and IL-6 were remarkably reduced in CIA mice treated with IL-1Ra-mRNA-LNP or IL-1Ra/ANL3-mRNA-LNP (Fig. 4B). These two cytokines also showed a trend of decrease in ANL3-mRNA-LNP treated mice. Frequencies of IFN-γ-or IL-4-secreting T cells were also analyzed. Interestingly, the IL-1Ra-mRNA-LNP monotherapy promoted the secretion of both IFN-γ and IL-4 from splenic CD4^+^ T cells, while this effect was minor in other treatment groups (fig. S3). The exact role of IFN-γ in RA development yet remains complex and controversial, which may differ depending on the disease stage. We further investigated on the changes of major T cell and B cell populations. Transcriptomic analysis of joint tissues revealed a slight increase in proportion of regulatory T (Treg) cells, suggesting a local immunosuppressive microenvironment induced by mRNA therapy (fig. S4A). In line with this, systemic analysis of the spleens showed an increasing trend of Treg cells, with bulk B cell and CD4^+^ T cell populations remaining largely unchanged (fig. S4B). Furthermore, expression of activation markers including PD-1 and ICOS on CD4^+^ T cells was unaltered across all treatments (fig. S4C).

**Figure 4.**
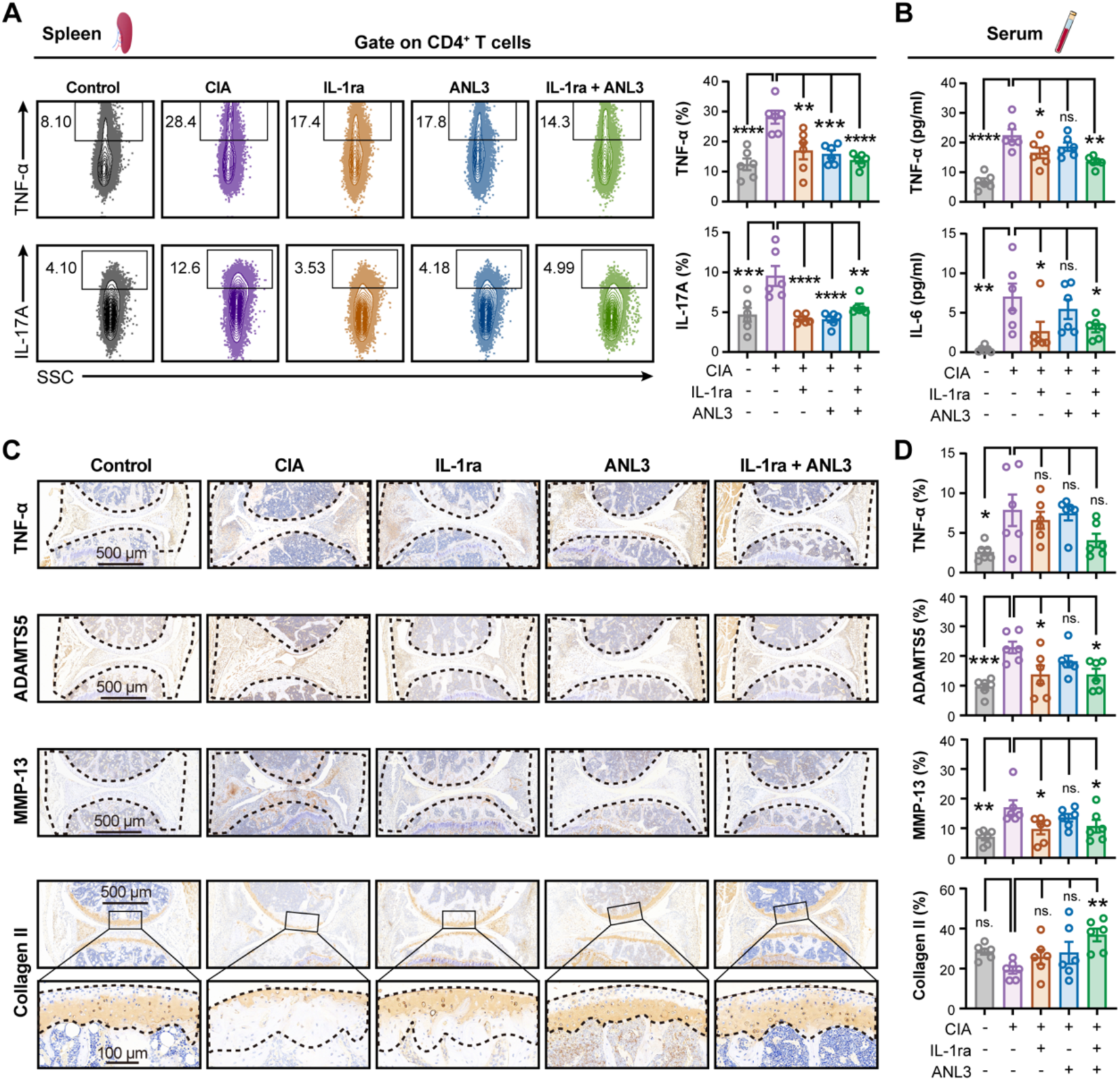
IL-1Ra/ANL3-mRNA-LNP therapy inhibits inflammation and promotes cartilage regeneration in CIA mice. **A,** Frequencies of TNF-α- or IL-17A-producing CD4^+^ T cells in spleens were measured by flow cytometric analysis (n=6/group)**. B,** Levels of serum TNF-α and IL-6 were analyzed by multiplexed cytometric bead array (n=6/group)**. C,** Immunohistochemical analysis of the indicated markers. Representative images of TNF-α, ADAMTS5, MMP-13 and Collagen II staining in the knee joints post treatment. **D,** Quantitative analysis of IHC-positive areas using Image J (n=6/group). Data are indicated as the mean ± SEM. Statistical significance was determined using One-way ANOVA with Tukey’s post hoc test. * *p* < 0.05; ** *p* < 0.01; *** *p* < 0.001; **** *p* < 0.0001.

We further examined the in-situ expression of TNF-α and representative factors regulating cartilage degradation and regeneration at the local knee joint sites. Although serum level of TNF-α and frequency of TNF-α-producing T cells were largely decreased by IL-1Ra-mRNA-LNP and IL-1Ra/ANL3-mRNA-LNP therapy (Fig. 4, A and B), TNF-α expression in synovial and local cartilage regions was only downregulated upon the combinatorial mRNA therapy (Fig. 4, C and D). IL-1Ra-mRNA-LNP monotherapy and the dual therapy efficiently suppressed expression of the matrix-degrading proteins ADAMTS5 and MMP-13 (Fig. 4, C and D). Most importantly, the combined treatment not only reversed collagen degradation but also robustly promoted collagen synthesis within the cartilage compartment that indicated tissue regeneration (Fig. 4, C and D). Collectively, these findings showed that the combinatorial mRNA therapy exerts a potent therapeutic efficacy by suppressing local and systemic inflammation and promoting cartilage regeneration simultaneously.

### IL-1Ra/ANL3-mRNA-LNP shows robust therapeutic efficacy in spontaneous RA model

Therapeutic potential of mRNA therapy was further evaluated in a TNF-α transgenic (TNF-Tg) mouse model over-expressing human TNF-α, which develops progressive and erosive arthritis spontaneously. In this regard, TNF-Tg mice with established arthritis at 14 weeks of age were intra-articularly administered with three doses of IL-1Ra-mRNA-LNP (2 μg), ANL3-mRNA-LNP (2 μg), or dual therapy with IL-1Ra/ANL3-mRNA-LNP (4 μg) at an interval of seven days (Fig. 5A). By week 18, mice from all treatment groups showed significant amelioration of arthritis symptoms, with the combinatorial therapy exhibiting the most pronounced effect. This was evidenced by a reduced arthritis index, diminished bone erosion, as well as alleviated degree of joint swelling (Fig. 5, B and D). Furthermore, histological analysis revealed that all treated mice, irrespective of receiving monotherapy or dual therapy, demonstrated a significantly reduced infiltration of inflammatory cells at the knee joint compartment (Fig. 5, E and F). In addition, cartilage thickness was markedly increased following mRNA treatment, as revealed by SO/FG and TBO staining (Fig. 5E). These data indicated that IL-1Ra/ANL3-mRNA-LNP treatment could efficiently mitigate the pathological manifestations of arthritis in TNF-Tg mouse model, and the therapeutic potency was superior to IL-1Ra-mRNA-LNP or ANL3-mRNA-LNP monotherapy.

**Figure 5.**
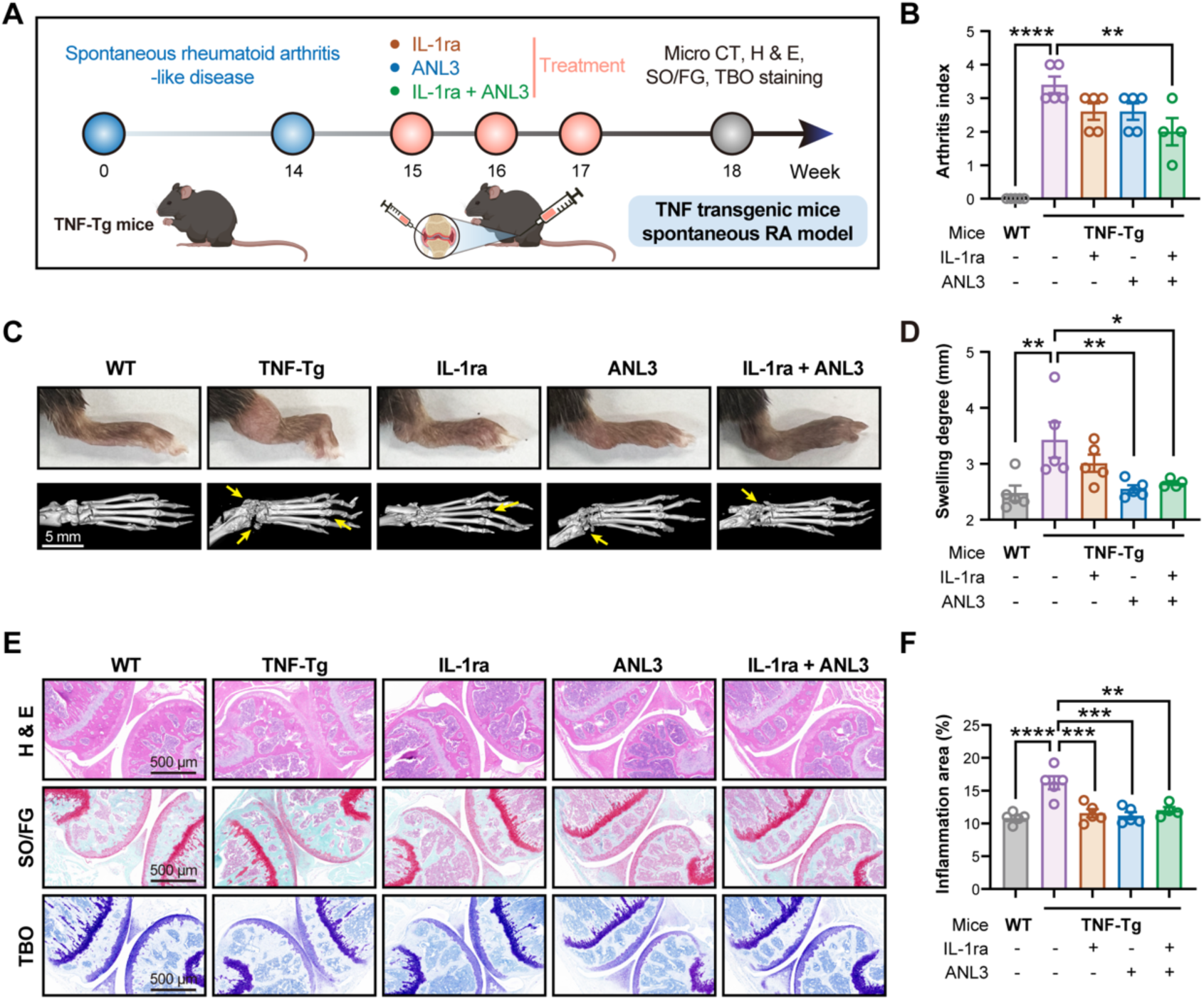
Therapeutic effect of mRNA therapies in TNF-Tg mice with spontaneous RA. **A,** Experimental design schematic. TNF-Tg mice spontaneously develop RA by week 14. From weeks 15 to 17, TNF-Tg mice received intra-articular administration of PBS (n=5), 2 μg of IL-1Ra-mRNA-LNP (n=5), 2 μg of ANL3-mRNA-LNP (n=5), or 4 μg of IL-1Ra/ANL3-mRNA-LNP (n=4) once a week for consecutive three weeks. Wild-type C57BL/6 mice were used as control (n=5). **B,** Arthritis index is shown. **C,** Representative images from morphological appearance (upper panel) and micro-CT scanning (lower panel) of ankle joints and paws of mice. Arrows indicate sites of bone erosion. **D,** Paw thickness indicative of swelling degree. **E,** Representative images of H&E, SO/FG and TBO staining of the tissue sections from knee joints. **F,** Inflammatory cell infiltration was quantified by measuring the percentage of inflammation-positive area using Image J. Data are displayed as the mean ± SEM. Statistical significance was determined using One-way ANOVA with Tukey’s post hoc test. * *p* < 0.05; ** *p* < 0.01; *** *p* < 0.001; **** *p* < 0.0001.

### IL-1Ra/ANL3-mRNA-LNP ameliorated OA-induced joint damage and regenerated cartilage

Given that OA and RA share significant similarities in terms of cartilage damage and joint inflammation. We further evaluated the potential of IL-1Ra/ANL3-mRNA-LNP in treating OA using DMM-induced mouse model, which is a commonly used surgical procedure in animal models for induction of OA that can mimic post-traumatic OA in humans. In this study, we established early-stage and late-stage OA models that differ in the time point of treatment initiation. Two-dose and three-dose treatment regimens were used for treating early-stage OA and late-stage OA, respectively (Fig. 6A). In the early-stage OA model, IL-1Ra/ANL3-mRNA-LNP significantly attenuated the cartilage damage, examined by H&E and SO/FG staining (Fig. 6B). Mankin’s and OARSI grading systems were further used to examine disease severity based on the histopathological features(*41*). It was found that the mRNA-treated mice had a remarkably lower Mankin’s score and OARSI score compared to the untreated mice (Fig. 6C). Immunohistochemical staining showed that mRNA treatment significantly reduced expression of matrix-degrading proteins ADAMTS5 and MMP-13 at the cartilage sites, and meanwhile promoted the local production of collagen II that was indicative of tissue regeneration (Fig. 6, D and E). We further performed the same analyses in late-stage OA mouse model and examined these key parameters. Following three doses of IL-1Ra/ANL3-mRNA-LNP treatment, cartilage damage was largely halted (Fig. 6, F and G), accompanied by a lower expression of ADAMTS5 and MMP-13 locally (Fig. 6, H and I). The treatment also increased production of collagen II, but the effect was less prominent than that observed in early-stage OA mice. We also evaluated the therapeutic potential of IL-1Ra/ANL3-mRNA-LNP in a severe OA model that started treatment 10 weeks post DMM surgery, while the treatment did not show adequate protection for cartilage damage (fig. S5).

**Figure 6.**
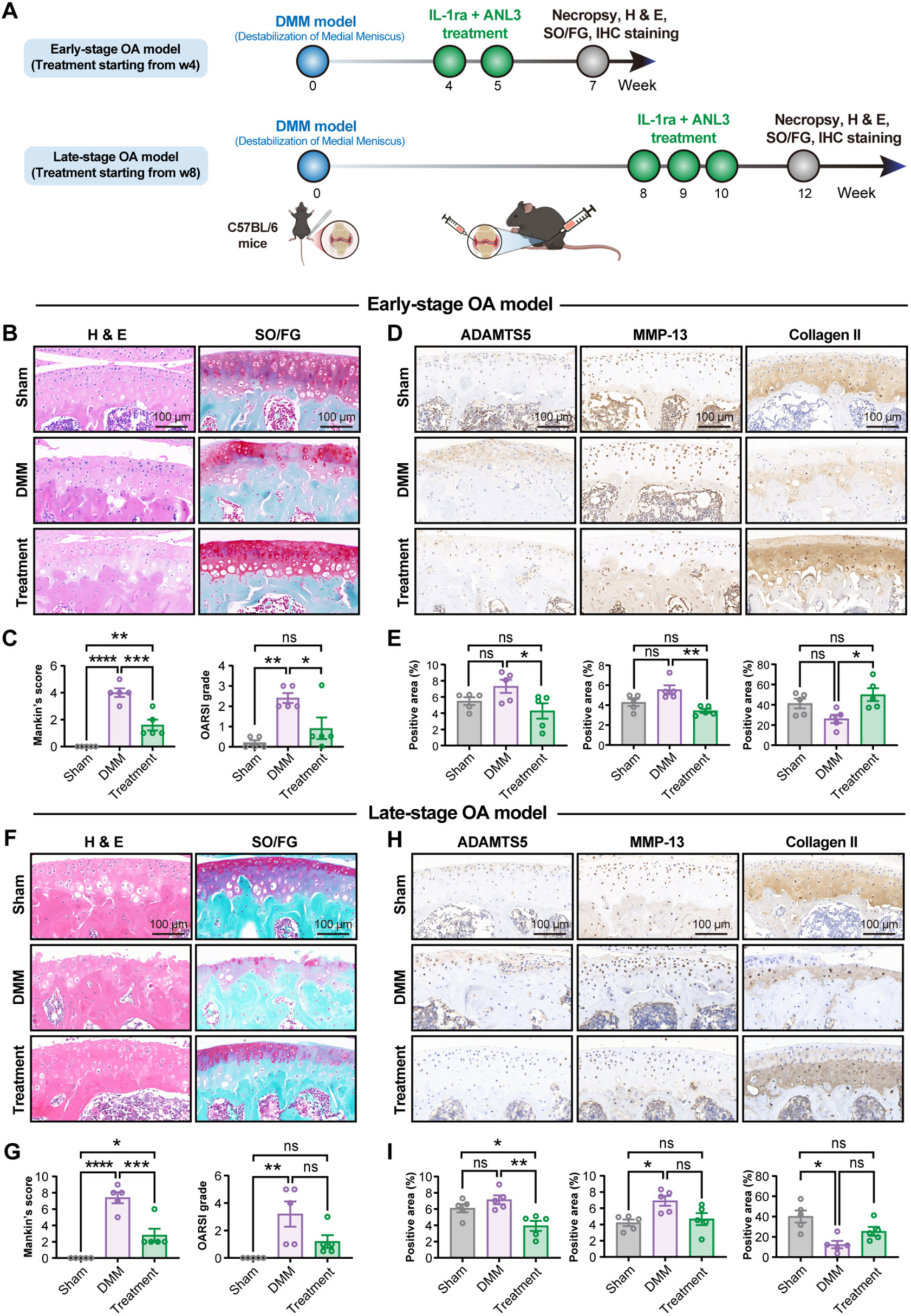
Therapeutic effect of mRNA therapies in DMM-induced surgical OA mice. **A,** Experimental design schematic. Early-stage OA mice and late-stage OA mice received intra-articular injections with two doses and three doses of IL-1Ra/ANL3-mRNA-LNP (4 μg), respectively. The sham-operated group represents healthy control. DMM mice treated with PBS serves as disease control (n=5/group). **B and F,** Representative images of H&E and SO/FG staining of the knee joints. **C and G,** Mankin’s score and OARSI grades are shown. **D and H,** Immunohistochemical analysis of the indicated markers. Representative images of ADAMTS5, MMP-13 and Collagen II staining in the knee joints post treatment. **E and I,** Quantitative analysis of IHC-positive areas from panel d and panel h, respectively. Data are shown as the mean ± SEM. Statistical significance was determined using One-way ANOVA with Tukey’s post hoc test. * *p* < 0.05; ** *p* < 0.01; *** *p* < 0.001; **** *p* < 0.0001.

### IL-1Ra/ANL3-mRNA-LNP demonstrated a high safety profile in mice

Finally, safety aspects of IL-1Ra/ANL3-mRNA-LNP was preliminarily assessed. Two chondrocyte cell lines, SW1353 and ATDC5, were incubated with escalating concentrations of mRNA drugs for 24 hours. Cell viability remained largely stable and was only marginally decreased upon the high-dose treatment, which suggested a very limited cytotoxicity in vitro (Fig. 7A). Further, alongside with the evaluation of therapeutic efficacy in late-stage OA mice (Fig. 6A), body weight was monitored upon the first treatment at week 8 and remained stable during the following four weeks of observation (Fig. 7B). At week 12, levels of serum alanine transaminase (ALT) and aspartate aminotransferase (AST) were both within the normal range without any increase post treatment (Fig. 7C). Specimens of main organs including heart, liver, lung and kidney were subjected to histopathological examination. H&E staining results revealed no pathological alterations in any of the groups (Fig. 7D). Safety profile of IL-1Ra/ANL3-mRNA-LNP was further validated in TNF-Tg mice alongside with the experiment described early (Fig. 5). At week 18, no changes were observed in terms of serum ALT, AST, Urea, and Creatinine, as well as the pathology of liver and kidney tissues (Fig. 7, E and F). These data altogether demonstrated that IL-1Ra/ANL3-mRNA-LNP was well-tolerated in arthritis mice showing a high-level safety profile.

**Figure 7.**
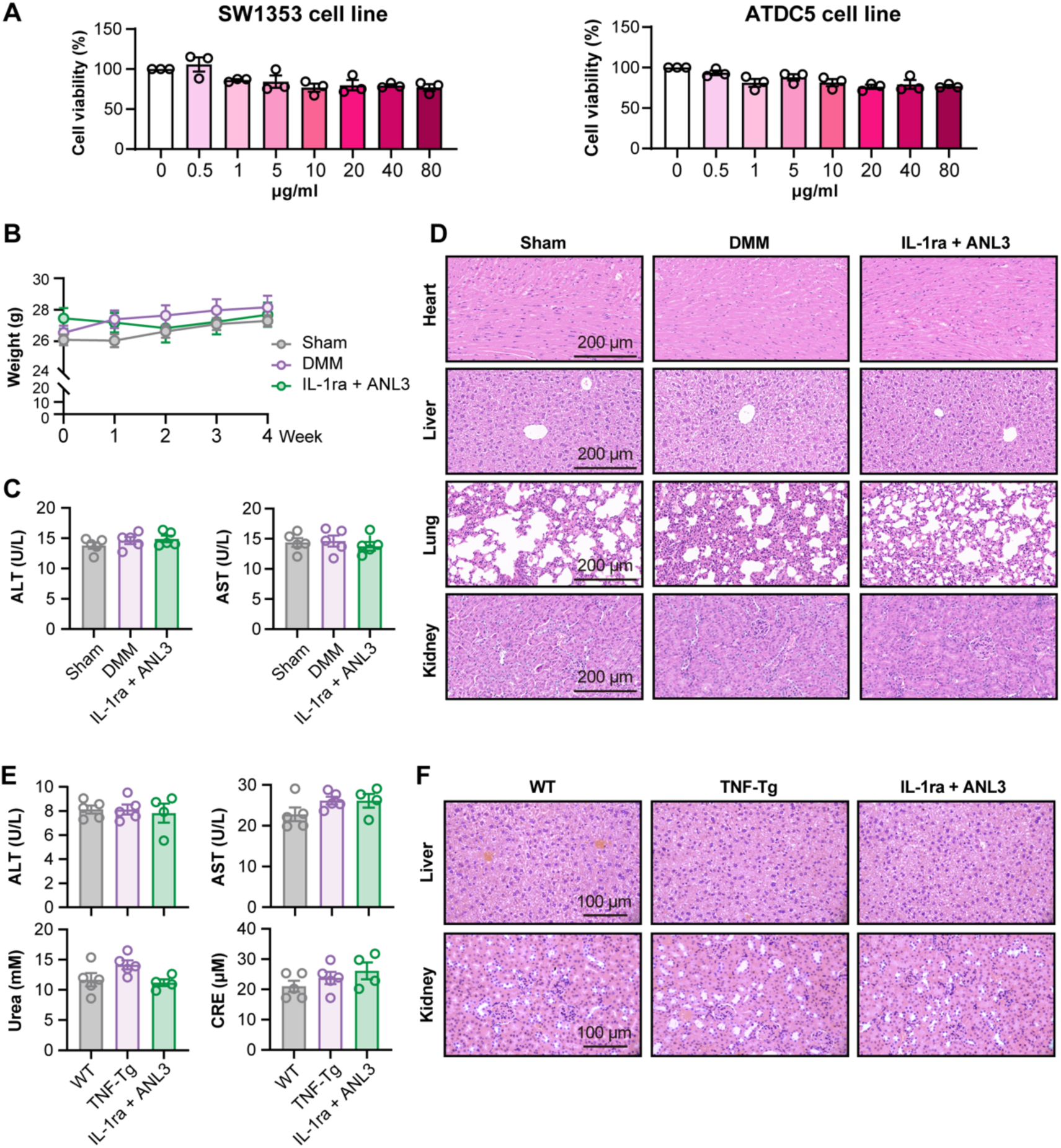
Preliminary evaluation of safety profile of IL-1Ra-ANL3-mRNA-LNP. **A,** Cell viability of human (SW1353) and mouse (ATDC5) chondrocyte cell lines following treatment with escalating doses of mRNA drug for 24 hours. Data from 3 independent experiments are show. **B-D,** Late-stage OA mice underwent treatment schedule shown in the Figure 6. Body weight of mice was monitored (**B**). Serum AST and ALT levels at the day of necropsy were assessed (**C**). Representative images of H&E sections of heart, liver, lung and kidney (**D**). **E-F,** TNF-Tg mice underwent treatment schedule shown in the Figure 5. Levels of serum ALT, AST, Urea and CRE were measured at week 18 (**E**). Representative images of H&E sections of liver and kidney (**F**). Data are indicated as the mean ± SEM.

## Discussion

RA and OA are two of the most prevalent and debilitating joint disorders, characterized by chronic pain, inflammation, and progressive destruction of articular cartilage. Despite their distinct etiologies, these two conditions converge on similar pathological outcomes involving inflammatory cascades, extracellular matrix degradation, and loss of cartilage integrity(*42*). However, current therapeutic options are largely palliative and fail to achieve desired effect. In this study, we designed a unique cartilage-penetrating LNP that delivers mRNAs encoding two complementary therapeutic proteins, IL-1Ra to antagonize inflammation and a pro-regenerative ANGPTL3 derivative (ANL3) to stimulate cartilage repair. This “one-shot” combinatorial therapy aims to treat arthritis by providing both an anti-inflammatory biologic and a regenerative factor together in a single administration.

Effective arthritis treatment should be able to suppress inflammation since chronic synovial inflammation drives cartilage matrix degradation via cytokines such as IL-1β and TNF-α(*43*). During the past decades, biologic DMARDs, for example Anakinra (recombinant IL-1Ra), has been approved but its therapeutic window was narrow, and duration of benefit was limited, which was largely due to its short intra-articular half-life and poor cartilage-penetrating capacity(*44*). To overcome this, we employed a unique mRNA-LNP platform to construct the IL-1Ra-mRNA modality for protein replacement therapy. A notable innovation in this study lies in the design of LNP with small size (60–70 nm) that can fit within the cartilage mesh for deeper penetration, allowing the encapsulated mRNA cargo to reach the chondrocytes in situ. This effect was mainly contributed by our proprietary ionizable cationic lipid FS01 that is featured by a squaramide headgroup and an ortho-butylphenyl-modified hydrophobic tail that facilitates lipid-mRNA interaction(*17*). Moreover, FS01 is naturally endowed with a low immunogenicity that can reduce inflammation driven by the lipid-based delivery vehicle itself(*17, 32*). Indeed, our comparative evaluation showed that FS01-LNP (∼60 nm) achieved superior intra-cartilage mRNA translation in vivo than three conventional LNPs (∼100 nm). By immunohistochemical approach, we dissected the location of IL-1Ra expression and found that it was precisely localized at the cartilage sites, which further validated its design rationale. We also noticed that a single intra-articular injection of IL-1Ra-mRNA yielded a significantly higher level of local protein expression than the counterpart recombinant IL-1Ra protein. This implies a key advantage of using mRNA-LNP for protein replacement therapy through a potential depot effect that may obviate frequent dosing. While when it comes to this point, it should also be noted that the mRNA used in our study was well designed through codon optimization, m1Ψ modification, as well as construction with a novel Cap 1 analog that collectively contribute to an efficient and sustained protein expression. However, the mRNA payload could be further optimized to achieve more durable expression, for example, by employment of the emerging circular mRNA (circRNA) technology.

Strategies to block inflammation (e.g. anti-IL-1 therapy) can only slow cartilage damage but fail to regenerate damaged tissue. This inspired us to include the modified form of ANL3, a well validated pro-regenerative factor, in our drug formulation aiming to promote chondrogenesis and restore joint homeostasis. The ANL3-mRNA was co-encapsulated with IL-1Ra-mRNA into LNP to formulate the drug. Through comprehensive evaluations in CIA mice, TNF-Tg mice and DMM surgical OA mice, this dual mRNA therapy demonstrated strikingly high potency in reducing joint swelling, bone erosion and synovial inflammation, outperforming the counterpart single-component mRNA therapy. These outcomes therefore reflected an effective IL-1 blockade by IL-1Ra-mRNA component. Moreover, in both RA and OA models, mice that received the combinatorial therapy retained a more uniform cartilage layer and increased cartilage thickness accompanied by robust expression of collagen II in situ compared to the untreated mice, which are direct evidence of anabolic repair and tissue regeneration. At the molecular level, transcriptomic profiling of treated joints provided further mechanistic validation. Dual mRNA therapy downregulated pathways related to IL-1, IL-6, and NF-κB signaling while concurrently suppressing extracellular matrix degradation pathways. Meanwhile, anabolic gene modules associated with chondrogenesis were largely upregulated. These data further confirmed a synergistic action that IL-1Ra primarily attenuates inflammation, whereas ANL3 reactivates cartilage matrix synthesis, together shifting the joint microenvironment from a catabolic, inflamed state towards an anabolic, reparative state.

Despite these promising results, several limitations of this study await further investigation. One critical point is the uncertainty of translation from mice to humans. Mouse joints are small and subject to rapid cartilage turnover, whereas human joints are much larger and may require higher doses or more frequent injections. Although preliminary safety assessments of the mRNA therapy demonstrated excellent tolerability in arthritis mice. RA and OA treatment normally requires repeated administrations, raising potential concerns about cumulative immune responses to mRNA-LNP components. Future studies should also assess durability and reversibility of the induced regenerative effects, as well as to further optimize this approach for prolonged protein expression to achieve more sustained efficacy.

In summary, this study establishes a versatile and rationally engineered dual mRNA therapy for treating RA and OA that combines two complementary mechanisms, anti-inflammatory and pro-regenerative. By overcoming the delivery and pharmacokinetic barriers of traditional biologics, this approach achieves sustained protein expression, potent disease suppression, and cartilage restoration in pre-clinical models of RA and OA. To our knowledge, no current arthritis therapy so far provides both an anti-inflammatory biologic and a regenerative factor together in one single administration. Our findings not only provide mechanistic insights into the coordinated control of inflammation and tissue repair but also illustrate the transformative potential of mRNA therapeutic for protein replacement therapy. With further optimization and validation in large-animal and human studies, the IL-1Ra/ANL3-mRNA-LNP therapy may represent a new class of disease-modifying immunomodulatory therapy capable of addressing the long-standing unmet needs in arthritis treatment.

## Acknowledgements

This work was supported by the National Natural Science Foundation of China (Grant nr. 32471004 to A.L., 82101896 to G.W., 82271839 to X.L. and 82401842 to Y.H.), Innovation and Entrepreneurship (ShuangChuang) Program of Jiangsu Province (Grant nr. JSSCTD202447 to A.L.), Jiangsu Province Graduate Research and Practical Innovation Program (Grant nr. KYCX24-1046 to K.G.), the Liaoning Provincial Education Department Basic Research Project (Grant nr. LJ212410161057 to G.W. and Grant nr. LJ212410161034 to X.L.), the Liaoning Provincial Science and Technology Department Joint Project (Grant nr. 2024-MSLH-113 to G.W.) and Dalian Medical University Interdisciplinary Research Cooperation Project Team Funding (Grant nr. JCHZ2023010 to X.L.). We thank the Targeted Discovery Center of China Pharmaceutical University for instrumental support on this study. We also thank Guangzhou Henovcom Bioscience for providing the proprietary LZCap agent for this study. We express our gratitude to online research tools, including OmicStudio and STRING. Additionally, we acknowledge the valuable contribution of BioRender in generating schematic diagrams.

## Author contributions

A.L., G.W., L.X. and Y.H. designed the study. Y.G., K.G., C.Z., Y.H., Y.Z., G.W. J.X., J.L., J.R., A.M., K.T., M.S. and X.C. performed the experiments and analysis. A.L., X.L., G.W., Y.Y., M.S., Y.H. discussed the data. A.L., G.W., Y.H. and K.G. wrote the manuscript. All authors have read and approved the manuscript.

## Data availability

All data are available upon reasonable request to the corresponding author. All raw sequencing data have been deposited in the open OMIX platform, the National Genomics Data Center (https://ngdc.cncb.ac.cn/omix; accession nr. OMIX012724).

