## Supplemental Materials for "A Combinatorial mRNA Therapy for Treating Rheumatoid Arthritis and Osteoarthritis by Inhibiting Inflammation and Promoting Cartilage Regeneration"

### Supplemental Materials and Methods

#### Cell line and cell culture

Human embryonic kidney 293T (HEK-293T) cells were purchased from Cell Resource Center, Shanghai Institutes for Biological Sciences, Chinese Academy of Sciences. Human chondrocytic cell line SW1353 was obtained from Haixing Biosciences (Suzhou, China). Mouse chondrocytic cell line ATDC5 was obtained from Cellverse Co., Ltd. (Shanghai, China). HEK-293T and ATDC5 cells were maintained in DMEM (BIOIND, Israel) containing 10% fetal bovine serum (FBS, BIOIND, Israel) and 1% penicillin-streptomycin (NCM Biotech, China). The SW1353 cell line was cultured in L-15 medium (HyCyte, China) supplemented with 10% fetal bovine serum and 1% penicillin-streptomycin. All cells were tested for mycoplasma contamination before use.

#### Cell viability assay

SW1353 and ATDC5 cells were tested for cell viability following LNP-mRNA treatment using cell counting kit-8 (CCK-8) reagent (SparkJade, China). Cells were seeded into 96-well plates at a density of 3000 cells/well. Upon adherence to plates for 24 hours, cells were treated with escalating doses of LNP-mRNA for 24 hours. After removal of medium, 100 μL of 10% CCK-8 solution was added to each well and incubated at 37 °C away from light for 1 hour. The absorbance was measured at 450 nm using a microplate reader (Thermo Scientific, USA).

#### Western blot assay

To evaluate the translation of ANL3-mRNA in vitro, HEK-293T cells were seeded into a 6-well plate and transfected with ANL3-mRNA using jetMESSENGER® transfection reagent according to the manufacturer's instruction. Following 48 hours of incubation, cells were lysed with RIPA buffer (Beyotime, China), and cell lysates were collected upon centrifugation (12,000 rpm, 15 min, 4 °C). Protein concentration was determined using a BCA Protein Quantification Kit (Vazyme, China). 20 μg of proteins were resolved via 10% SDS-PAGE and transferred onto PVDF membranes (Immobilon, Millipore). Membranes were blocked with 5% non-fat milk in TBST (TBS containing 0.075% Tween-20) for 1 hour at room temperature (RT), followed by incubation with anti-GAPDH Ab (1:5000, Proteintech) or anti-ANGPTL3 Ab (1:5000, Abcam) overnight at 4 °C. After washing steps, membranes were incubated with HRP-conjugated anti-mouse IgG (1: 5000, Proteintech) for 2 hours at RT. The membrane was washed, and signal was detected with the ECL substrate (Vazyme, China) on a Tanon 5200 Multi (Shanghai, China).

**Enzyme Linked Immunosorbent Assay (ELISA)**

To evaluate the translation of IL-1Ra-mRNA in vitro, IL-1Ra-mRNA (2 μg) was transfected into HEK-293T cells. Cell culture supernatants were collected upon centrifugation (1500 rpm, 5 min, 4 °C). Cell lysates were collected upon centrifugation (12,000 rpm, 15 min, 4 ℃). Protein concentration was determined using a BCA Protein Quantification Kit (Vazyme, China). After this, IL-1Ra protein levels in culture supernatants and cell lysates were measured using a Mouse IL-1Ra/IL-1F3 ELISA Kit (MULTI SCIENCES, China) according to the instruction.

**Histological analysis and immunohistochemical staining**

For histological analysis, mouse knee joints were fixed in 4% paraformaldehyde overnight, decalcified in 14% EDTA, and embedded in paraffin. 5-μm tissue sections were cut, dewaxed and rehydrated through xylene and alcohols and were subjected to staining with H&E, SO/FG, or Toluidine Blue. For the experiments evaluating safety, the heart, liver, lung, and kidney specimens were fixed with 4% paraformaldehyde, sectioned, and evaluated by H&E staining for pathological evaluation. For Immunohistochemical (IHC) analysis, tissue sections were dewaxed, rehydrated and antigen retrieved using proteinase K antigen retrieval solution (Abcam). Following this, an appropriate amount of endogenous peroxidase blocker was applied at RT for 10 minutes. The sections were then blocked with 5% BSA at RT for 30 minutes, followed by incubation overnight at 4°C with primary antibodies including anti-mouse TNF-α (1:200, Servicebio, China), anti-mouse ADAMTS5 (1:100, Affinity, USA), anti-mouse MMP-13 (1:100, Servicebio, China), or anti-mouse type II collagen (1:200, Servicebio, China) in a humidified chamber. A universal two-step detection kit (Servicebio, China) was used according to the manufacturer's instructions for secondary antibody incubation, followed by DAB chromogenic staining and hematoxylin counterstaining. Finally, the sections were dehydrated through a graded ethanol series, cleared in xylene, and mounted with neutral resin. Images were captured using a digital pathological slide scanner (LG-S80, Servicebio, China).

**Preparation of single cell suspension**

Spleen tissues were gently grounded and passed through a 40-μm sterile cell strainer. Cells were then suspended in PBS and centrifuged at 400 g for 10 min. To disrupt red blood cells (RBCs), cell pellets were resuspended with RBC lysis buffer (Solarbio) for 5 min at 4 °C. Subsequently, 1× PBS was used to halt the lysis process, and the cells were washed at 400 g for 10 min to obtain splenic mononuclear cells. These cells were then re-suspended in RPMI-

1640 medium supplemented with 10% FBS (BIOIND) and 1% penicillin-streptomycin (NCM Biotech) for further experiments.

**Multiplexed cytometric bead array (CBA)**

Sera samples collected from mice were analyzed for specific cytokine expression. Inflammatory cytokines were quantified using a Mouse Inflammation CBA Kit (BD Biosciences, USA) according to the manuals. Samples were acquired in flow cytometer cytometer (FACS Fortessa, BD Biosciences, USA). Data were analyzed using FCAP Array Software Version 3.0 (BD, USA).

**RNA sequencing and Transcriptome analysis**

Total RNA was extracted from the knee joint tissues of DBA/1J mice (n=6/group) according to protocol reported previously[1]. Concentration and integrity of RNA were assessed using a Qubit fluorometer and Qsep400 high-throughput fragment analyzer. RNA samples from two mice per group were pooled together for sequencing. The RNA sequencing was performed at Metware Biotechnology Co., Ltd. (Wuhan, China). For library construction, mRNA was first enriched using oligo (dT) magnetic beads and the purified mRNA was further fragmented into short pieces using fragmentation buffer under appropriate temperature conditions. 2 μg of total RNAs was used for preparation of the stranded RNA sequencing library. The library products (250-350bps) were quantified using a Qubit fluorometer, and fragment size distribution was checked with the Qsep400 analyzer. Qualified libraries were then subjected to Illumina sequencing. Raw reads were filtered using fastp according to the following criteria: Reads containing adapter sequences were removed. Reads with more than 10% ambiguous bases (N) were discarded. Reads with over 50% low-quality bases ($Q \leq 20$) were removed. Subsequent analyses were all based on the resulting clean reads. Bioinformatic analysis (including principal component analysis, differential analysis, enrichment analysis, and immune infiltration analysis) was performed using the OmicStudio tools. The differentially expressed genes (DEGs) were defined as genes with a $p$-value $< 0.05$, $|\log2 \text{ fold change}| \geq 1$, and $FDR < 0.05$. Protein-protein interaction (PPI) network analysis was performed on the DEGs using interaction data from the STRING database (http://string-db.org)[2]. All raw sequencing data have been deposited in the open OMIX platform, the National Genomics Data Center (https://ngdc.cncb.ac.cn/omix; accession nr. OMIX012724).

**Determination of liver and renal function**

The concentrations of aspartate aminotransferase (AST), alanine aminotransferase (ALT), carbamide (Urea), and creatinine (CRE) were measured in serum using the Aspartate aminotransferase Assay Kit (Ruixinbio, China), Alanine aminotransferase Assay Kit (Ruixinbio, China), Urea Assay Kit (Ruixinbio, China) and Creatinine Assay Kit (Ruixinbio, China) according to the manufacturer's instructions, respectively.

**Supplementary Figure 1.**

| Ionizable lipids | Lipid 5 | SM102 | FS01 | LP01 |
|---|---|---|---|---|
| Molecular weight | 710.17 | 710.2 | 865.69 | 852.27 |
| pKa | 6.56 | 710.2 | 6.45 | 6.1 |
| Particle size (nm) | 87.24 ± 1.25 | 87.22 ± 2.06 | 67.06 ± 1.67 | 90.65 ± 2.33 |
| Polydispersity Index | 0.134 ± 0.05 | 0.114 ± 0.03 | 0.156 ± 0.022 | 0.144 ± 0.032 |
| Zeta potential (mV) | -3.37 ± 0.49 | -1.57 ± 4.92 | 2.24 ± 0.878 | 2.68 ± 2.238 |
| Encapsulation efficiency | 94.43 ± 0.80 | 95.63 ± 0.55 | 96.97 ± 1.66 | 92.57± 1.93 |

**Figure S1. Physiochemical parameters of ionizable lipids and their corresponding LNPs.** A summary of physicochemical properties of ionizable lipids and their corresponding LNPs, including molecular weight, pKa, particle size, polydispersity index (PDI), zeta potential, and encapsulation efficiency.

**Supplementary Figure 2.**

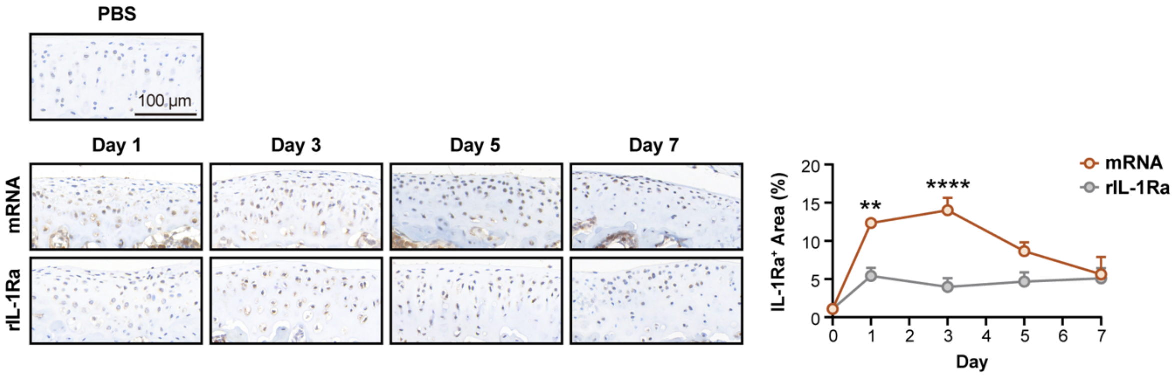


**Figure S2. Pharmacokinetics of IL-1Ra at cartilage sites upon intra-articular administration with IL-1Ra-mRNA-LNP or recombinant IL-1Ra.**

IL-1Ra-mRNA-LNP (2μg) and recombinant rIL-1Ra (2μg) were administered into C57BL/6 mice (n = 4 per group) via intra-articular injection. Knee joints were collected at the indicated time points, and immunohistochemical (IHC) staining was performed to quantify protein expression. Representative IHC images are shown (left), alongside corresponding quantitative analyses (right). Scale bar = 100 μm.

**Supplementary Figure 3.**

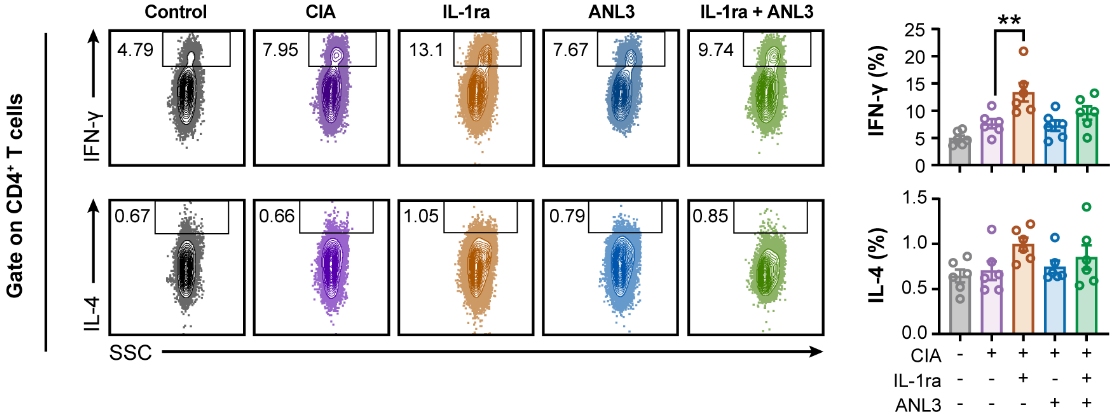


**Figure S3. Measurement of cytokine-producing CD4$^+$ T cells in spleens.**

Frequencies of IFN-γ- or IL-4-producing CD4$^+$ T cells in spleens of mice were measured by flow cytometric analysis (n=6/group). Data are indicated as the mean ± SEM. Statistical significance was determined using One-way ANOVA with Tukey's post hoc test. ** $p < 0.01$.

**Supplementary Figure 4.**

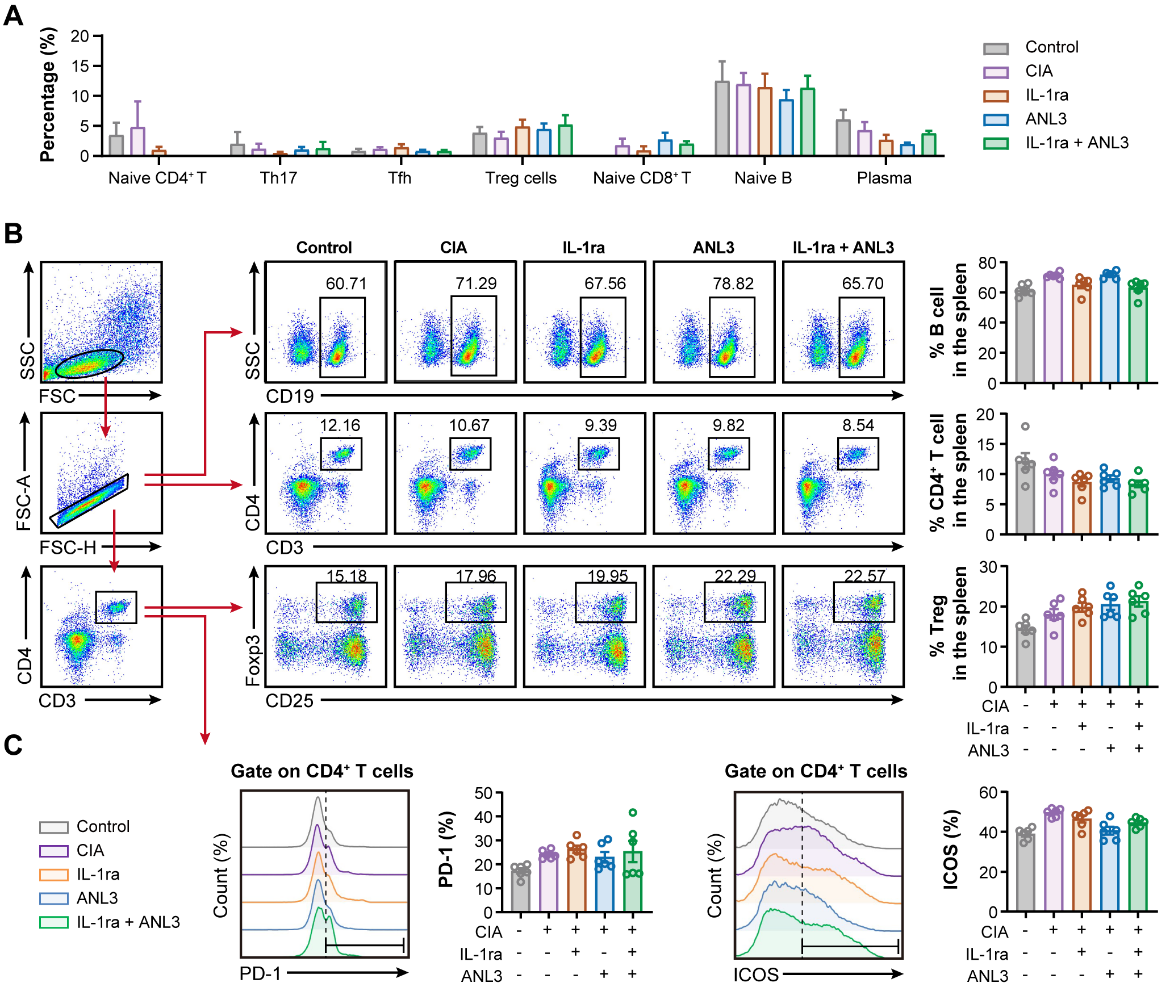


**Figure S4. Changes in frequencies of major T cell and B cell populations in CIA mice post mRNA-LNP treatment.**

CIA mice were treated following the experimental schedule shown in Figure 2b. At day 55, mice were necropsied, and knee joint specimens were collected for RNA-seq analysis. **a,** Immune infiltration analysis reveals the proportions of the indicated immune cell subsets within knee joint tissues (n=3/group). **b,** Frequencies of bulk B cells ($CD19^+$), $CD4^+$ T cells, and Treg cells ($CD3^+CD4^+CD25^+Foxp3^+$) in spleens were analyzed by flow cytometry (n=6/group). **c,** Surface expression of PD-1 and ICOS on splenic $CD4^+$ T cells were measured by flow cytometry (n=6/group). Data are expressed as the mean ± SEM.

**Supplementary Figure 5.**

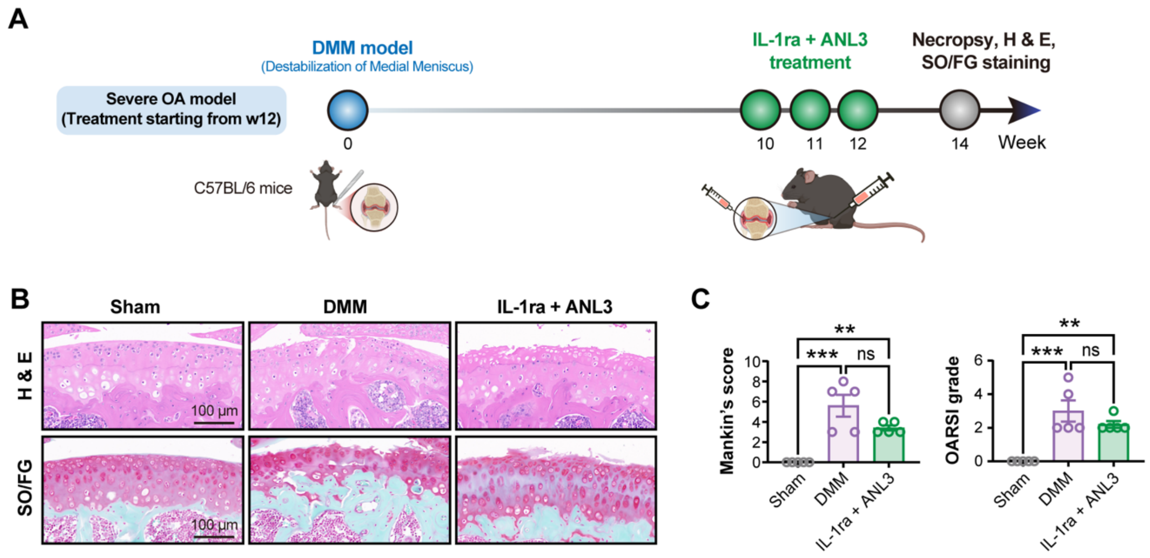


**Figure S5. Therapeutic effect of mRNA therapies in a severe OA model.**

**a,** Experimental design schematic. A severe OA mouse model was established and used. mRNA treatment starts at week 10 post DMM surgery. **b,** Representative images of H&E and SO/FG staining of the knee joints. **c,** Mankin's score and OARSI grades are shown. Data are shown as the mean ± SEM. Statistical significance was determined using One-way ANOVA with Tukey's post hoc test. ** $p < 0.01$; *** $p < 0.001$.

**Supplementary Table 1.**

Table S1. Sequences of IL-1Ra-mRNA and ANL3-mRNA

| Name | 5'-UTR | CDS region | 3'-UTR |
|---|---|---|---|
| **IL-1Ra** | AGAATAA<br>ACTAGTAT<br>TCTTCTGG<br>TCCCCAC<br>AGACTCA<br>GAGAGAA<br>CCCGCCA<br>CC | ATGGAGACCCCTGCTCAGCTGCTGTTCCT<br>GCTCCTGCTGTGGCTGCCTGACACCACCG<br>GAAGGCCTAGCGGCAAGAGGCCTTGCAA<br>GATGCAAGCCTTTAGGATCTGGGACACCA<br>ATCAGAAGACCTTCTACCTGAGGAACAAT<br>CAGCTGATCGCCGGCTACCTGCAAGGCCC<br>TAACATCAAGCTGGAGGAGAAGATCGACA<br>TGGTGCCTATCGACCTGCACAGCGTGTTC<br>CTGGGCATCCACGGCGGCAAGCTGTGCCT<br>GAGCTGCGCCAAGAGCGGCGACGACATC<br>AAGCTGCAGCTGGAAGAGGTGAACATCA<br>CCGACCTGAGCAAGAACAAGGAGGAGGA<br>CAAGAGGTTCACCTTCATTAGGAGCGAGA<br>AGGGCCCTACCACAAGCTTCGAGAGCGCC<br>GCCTGCCCTGGCTGGTTCCTGTGCACCAC<br>CCTGGAGGCCGATAGGCCTGTGAGCCTGA<br>CCAACACCCCTGAGGAGCCTCTGATCGTG<br>ACCAAGTTCTACTTCCAAGAGGATCAGTG<br>ATGA | CTGGTACTGCATGCA<br>CGCAATGCTAGCTGC<br>CCCTTTCCCGTCCTG<br>GGTACCCCGAGTCTC<br>CCCCGACCTCGGGTC<br>CCAGGTATGCTCCCA<br>CCTCCACCTGCCCCA<br>CTCACCACCTCTGCT<br>AGTTCCAGACACCTC<br>CCAAGCACGCAGCA<br>ATGCAGCTCAAAAC<br>GCTTAGCCTAGCCAC<br>ACCCCCACGGGAAA<br>CAGCAGTGATTAACC<br>TTTAGCAATAAACGA<br>AAGTTTAACTAAGCT<br>ATACTAACCCCAGGG<br>TTGGTCAATTTCGTG<br>CCAGCCACACC |
| **ANL3** | AGAATAA<br>ACTAGTAT<br>TCTTCTGG<br>TCCCCAC<br>AGACTCA<br>GAGAGAA<br>CCCGCCA<br>CC | ATGGAGACCCCCGCTCAGCTGCTGTTCCT<br>GCTCCTGCTGTGGCTGCCCGACACCACCG<br>GCATCCCCGCCGAGTGCACCACCATCTAC<br>AACAGAGGCGAGCACACAAGCGGCATGT<br>ACGCCATCAGACCTAGCAACAGCCAAGTG<br>TTCCACGTGTACTGCGACGTGATCAGCGG<br>CAGCCCCTGGACCCTGATTCAGCACAGAA<br>TCGACGGCAGCCAAAACTTCAACGAGAC<br>CTGGGAGAACTACAAGTACGGCTTCGGCA<br>GACTGGACGGCGAGTTCTGGCTGGGCCTG<br>GAGAAGATCTACAGCATCGTGAAGCAGAG<br>CAACTACGTGCTGAGAATCGAGCTGGAGG<br>ACTGGAAGGACAACAAGCACTACATCGA<br>GTACAGCTTCTACCTGGGCAACCACGAGA<br>CCAACTACACCCTGCACCTGGTGGCCATC<br>ACCGGCAACGTGCCCAACGCCATCCCCGA<br>GAACAAGGACCTGGTGTTCAGCACCTGGG<br>ACCACAAGGCCAAGGGCCACTTCAACTGC<br>CCCGAGGGCTACAGCGGCGGCTGGTGGTG<br>GCACGACGAGTGCGGCGAGAACAACCTG<br>AACGGCAAGTACAACAAGCCTAGAGCCA<br>AGAGCAAGCCCGAGAGACGGAGAGGCCT<br>GAGCTGGAAGAGCCAAAACGGCAGACTG<br>TACAGCATCAAGAGCACCAAGATGCTGAT<br>CCACCCCACCGACAGCGAGAGCTTCGAGT<br>GATGA | CTGGTACTGCATGCA<br>CGCAATGCTAGCTGC<br>CCCTTTCCCGTCCTG<br>GGTACCCCGAGTCTC<br>CCCCGACCTCGGGTC<br>CCAGGTATGCTCCCA<br>CCTCCACCTGCCCCA<br>CTCACCACCTCTGCT<br>AGTTCCAGACACCTC<br>CCAAGCACGCAGCA<br>ATGCAGCTCAAAAC<br>GCTTAGCCTAGCCAC<br>ACCCCCACGGGAAA<br>CAGCAGTGATTAACC<br>TTTAGCAATAAACGA<br>AAGTTTAACTAAGCT<br>ATACTAACCCCAGGG<br>TTGGTCAATTTCGTG<br>CCAGCCACACC |

**Supplementary Table 2.**

Table S2. A list of anti-mouse antibodies used for FACS analysis.

| Antibody | Clone | Manufacturer |
|---|---|---|
| Anti-Mouse CD3 | 145-2C11 | Biolegend |
| Anti-Mouse CD4 | GK1.5 | Biolegend |
| Anti-Mouse IFN-γ | XMG1.2 | Biolegend |
| Anti-Mouse IL-4 | 11B11 | eBioscience |
| Anti-Mouse TNF-α | MP6-XT22 | eBioscience |
| Anti-Mouse CD3 | 145-2C11 | eBioscience |
| Anti-Mouse CD25 | PC61 | Biolegend |
| Anti-Mouse FoxP3 | FJK-16S | eBioscience |
| Anti-Mouse ICOS | C398.4A | Biolegend |
| Anti-Mouse PD-1 | J43 | eBioscience |
| Anti-Mouse CD19 | 6D5 | Biolegend |
| Anti-Mouse IL-17A | eBio17b7 | eBioscience |
